# Saving Old Bones: a non-destructive method for bone collagen prescreening

**DOI:** 10.1101/653212

**Authors:** Matt Sponheimer, Christina M. Ryder, Helen Fewlass, Erin K. Smith, William J. Pestle, Sahra Talamo

## Abstract

Bone collagen is an important material for radiocarbon, paleodietary, and paleoproteomic analyses, but it degrades over time. Various methods have been employed to prescreen bone for collagen preservation, but these are often destructive and/or require exportation for analysis. Here we show that near-infrared spectroscopy can be used to determine bone collagen content quickly and non-destructively on site.r

## Introduction

The persistence of organic molecules in bone has proven crucial for understanding the human past. Bone collagen (a common protein in bone and skin) from humans and our close kin has been used to radiocarbon (^14^C) date crucial events in human history, such as the colonization of the Americas^1^ and southeastern Europe^2^ and the disappearance of groups including the Neanderthals^3,4^. In fact, much of our understanding of the sequence of human history prior to the advent of writing and calendrical systems comes from radiocarbon dating of bone collagen or other organic materials from archaeological sites. Bone collagen is also a preferred material for stable isotopic paleodietary studies and has been used to document the emergence of maize agriculture^5^, the broadening of European *Homo sapiens’* resource base in the Upper Paleolithic, and the importance of animal proteins in the diets of Neanderthals among other things^6^. It is also of increasing interest for paleoproteomic analyses, as it can be used to identify modern and ancient species and their phylogenetic histories even where ancient DNA studies are impossible or impractical^7^. Thus, it is fair to say that collagen is a material of signal importance for revealing the often murky human past, and that our ability to discern distant human behavior and evolutionary history will be roughly proportional to its preservation in the archaeological record.

Lamentably, collagen deteriorates over time, making it progressively more difficult to conduct analyses of these kinds as material gets older, although the speed of its degradation is heavily dependent on environmental conditions^8^. Moreover, preservation between and within individual archaeological or paleontological sites is highly variable. As a result, even recent sites may preserve little or no collagen, and ancient sites where collagen preservation is generally poor may have specimens that are surprisingly well preserved^9,10^. As a result, radiocarbon, paleodietary, and other archaeometric labs may need to destructively sample large numbers of specimens with the hope of finding a few suitable for analysis. This is not only ethically problematic, but it means that much time, effort, and money must go into the initial analysis and preparation of samples that will not yield meaningful results.

Consequently, there is intense interest in the development of methods to prescreen bone for collagen content while minimizing damage to specimens. Arguably, at least in the radiocarbon community, the standard method for determining a bone’s suitability for subsequent analysis is to take small subsamples (<5mg) for elemental analysis where %N, and to a lesser extent C/N ratios, are used to estimate collagen preservation^10,11^. In general, samples with more than 0.76% N by weight are considered likely to retain more than 1% collagen, which is typically sufficient for ^14^C analysis^10–12^. Other prescreening methods include mid-infrared or Raman spectroscopy which reveal information about a substance’s functional groups due to its interaction with electromagnetic radiation^13–18^. Although both elemental and spectroscopic techniques are clearly useful, they are often time-consuming, destructive, and/or typically require removal of bones from sites or museums to labs for analysis.

There have been, however, a few attempts to circumvent these limitations. Most notably, a portable Raman spectrometer with a 1060 nm laser was used to show that the ratio of peaks at 1450 cm^−1^ to 960 cm^−1^ is strongly associated with collagen content (R^2^=0.84)^13^. Additionally, qualitative non-destructive near-infrared (NIR) spectroscopy was used to classify 16 Holocene bones and four validation samples into poor and good collagen preservation groups^19^. Here, we build on the latter study to show that a portable and field ruggedized NIR spectrometer can be used not only to classify bones into groups by preservation status, but also to quantify percent collagen preservation (hereafter %coll) in ground and whole bone of Late Pleistocene age. NIR spectroscopy has great potential for bone prescreening in that it is non-destructive, has a very fast speed of analysis (typically seconds), is readily miniaturized so that field-deployable instruments are widely available, and has a greater effective penetration depth than its spectroscopic siblings (millimeters as opposed to microns; Supplementary Fig. 1). Its ability to provide a deeper glimpse is especially important given that bone surfaces are often heavily modified post-depositionally^14^.

## Results & Discussion

The near-infrared spectra of archaeological specimens with differing collagen contents are clearly distinct and multiple bands/regions, including the first overtone of the C-H stretch at 1700-1740 nm, N-H stretching combinations at 2050 nm, the N-H bend second overtone and C=O stretch combinations at 2180 nm, and C-H combinations at 2270-2290 nm show expected differences related to %coll (Figure 1a; see Methods)^20^. An exploratory principal component analysis (PCA) of the NIR spectra from 50 ground bone specimens from archaeological sites of Holocene age from the Old and New Worlds (Supplementary Table 1) reveals strong spectral differences between specimens with differing collagen contents (Figure 1b) and bands/regions associated with collagen (above) are highly influential in the loadings plot for PC1 and PC2 (Figure 1c).

**Figure 1:**
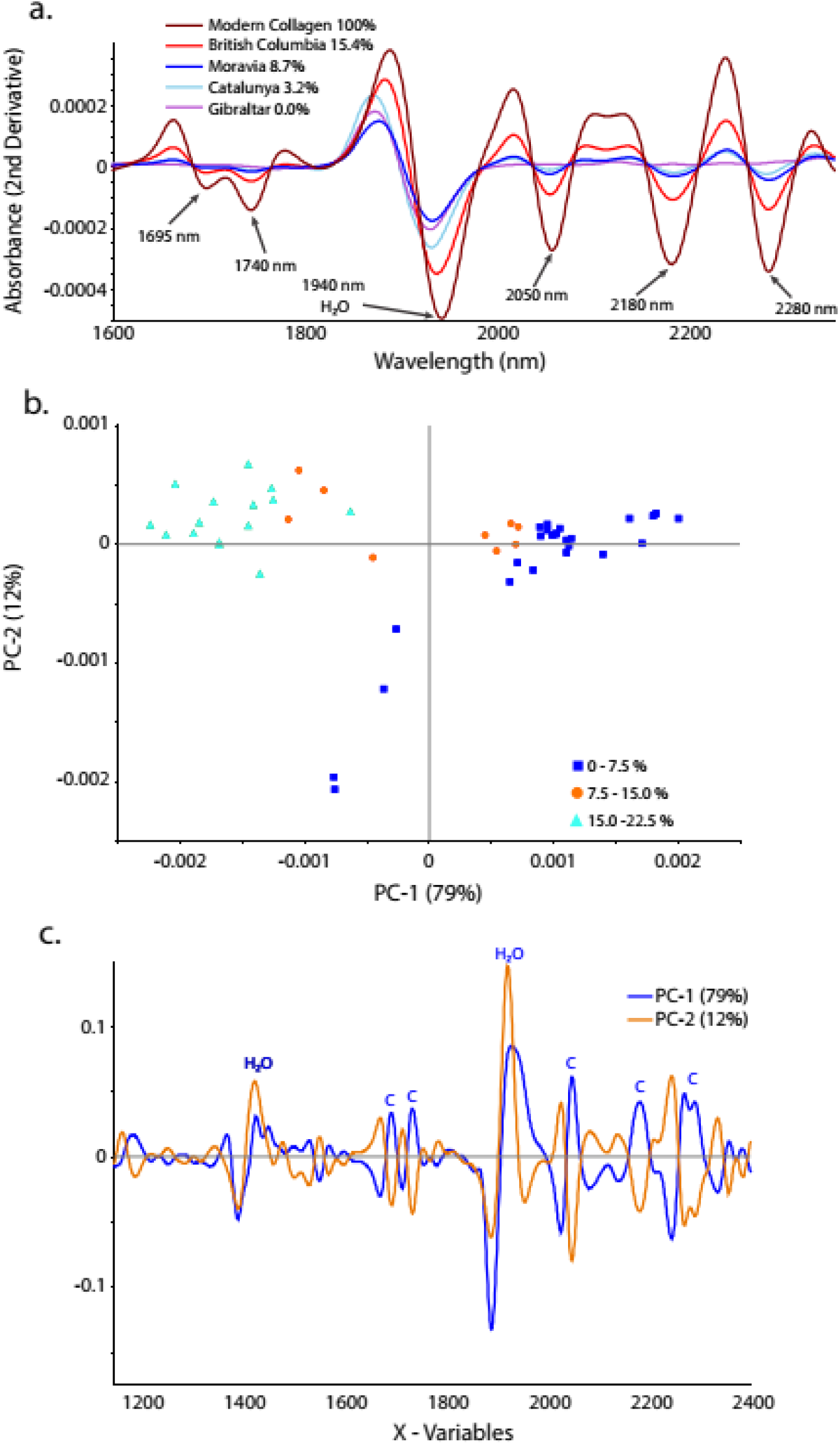
NIR bands reflect collagen content. A) Second derivative of near-infrared spectra from pure modern collagen (100% collagen; brown) and archaeological specimens from Gibraltar (0.0% collagen; pink), Catalunya (3.2% collagen; turquoise), Moravia (8.7% collagen; blue), and British Columbia (15.4% collagen; red). Multiple bands/regions show expected directional shifts in accordance with % collagen including the first overtone of the C-H stretch at 1695-1740 nm, N-H stretching combinations at 2050 nm, the N-H bend second overtone and C=O stretch combinations at 2180 nm, and C-H combinations at 2270-2290 nm. The region associated with water (and to a lesser extent protein) at around 1940 nm (O-H bend second overtone and O-H stretch + O-H deformation combination) was excluded from all analyses. B) PCA scores plot (PC1 and PC2) of the NIR spectra (780 nm to 2500 nm; second derivative) of 50 ground bone samples from archaeological sites. High collagen specimens (15.0% to 22.5%; turquoise triangles) are distinct from low collagen specimens (0% to 7.5%; blue squares) while samples with middling collagen contents (7.5% to 15%; orange circles) fall between these two groups. C) PCA loadings plot showing influential variables for PC1 (orange) and PC2 (blue). Bands/regions associated with collagen (labeled C; 1695-1740 nm, 2050 nm, 2180 nm, 2270-2290 nm) load on PC1 and PC2. Two regions associated with water (1430-1450 nm and 1940 nm) were kept out of subsequent analyses.

Partial least squares regression (PLSR) on a calibration dataset (25 spectra) resulted in a two factor model that predicted %coll from spectral data remarkably well (R^2^ = 0.98, Root-Mean Square Error of Calibration [RSMEC] = 1.00)(Figure 2a). The model also performed well when used to predict %coll from 25 independent validation specimens (R^2^=0.94; Root-Mean Square Error of Prediction [RSMEP] = 1.8)(Figure 2b). Of the 20 specimens in the validation sample with more than 3% collagen, the model predicted more than 3% collagen 20 times (100% classification success). The model correctly predicted all five specimens with less than 3% collagen (100% classification success).

**Figure 2:**
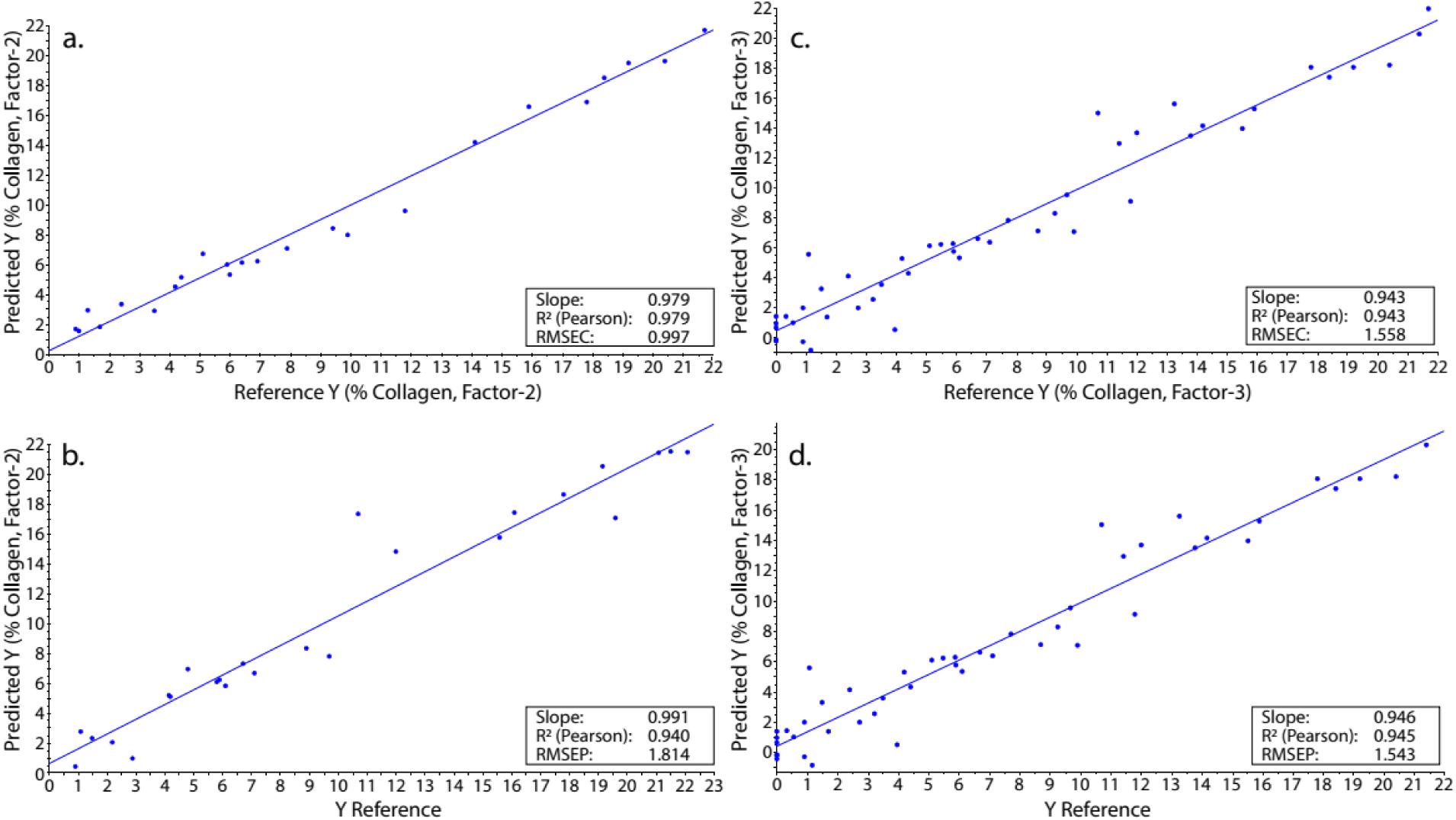
Predicting collagen preservation from NIR spectra. A) Results of PLSR showing predicted versus actual %coll values for the 25 sample (ground bone) calibration dataset (R2 = 0.98). B) Predicted versus actual %coll values using the calibration model on the 25 sample (ground bone) validation dataset (R2 = 0.94). C) Results of PLSR showing predicted versus actual %coll values for the 48 specimen ground/whole bone calibration dataset (R2 = 0.94). D) Predicted versus actual %coll values using the calibration model on the 49 sample validation dataset (R2 = 0.95).

Given the success of the model with ground bone samples, we built a model that included ground and whole bone samples and that included bone up to about 45 thousand years old (Supplementary Table 1). PLSR on spectra from a 48 sample calibration dataset performed very well (R^2^ = 0.94; RMSEC = 1.56; 3 factors; Figure 2c) and it predicted %coll in a 49 sample validation dataset equally well (R^2^=0.95; RMSEP = 1.54; Figure 2d). Of the 31 specimens in the validation dataset with more than 3% collagen, the model predicted 29 correctly (94% classification success). Of the 18 specimens with less than 3% collagen it predicted 84% correctly. Most crucially, of the 32 specimens in the validation dataset that the model predicted had more than 3% collagen, every one had more than the 1% collagen typically required for radiocarbon and paleodietary analyses.

These results demonstrate that NIR spectroscopy can be used to ascertain collagen preservation status in archaeological bone from dozens of sites across the world which range in age from recent to more than 45,000 years old. It is likely, therefore, that this tool will prove generalizable. Sites often contain large numbers of bones or fragments that might prove useful for radiocarbon, paleodietary, or paleoproteomic analyses. Even sites with relatively poor bone preservation (typically less than 1% collagen by weight) can contain specimens with reasonably intact collagen. For instance, of 50 bones analyzed from the Neanderthal site Zafarraya, six retained more than 4% collagen^10^. NIR spectroscopy might be used to quickly separate the wheat from the chaff at such sites.

Another benefit of this technique would be the ability to more adequately pick specific spots of individual bones for analysis. Archaeological bone is far from homogenous when it comes to collagen preservation^10^, yet typically only one small area is sampled to determine a bone’s suitability. It is easy to envision taking NIR scans of multiple spots on a bone (or taking hyperspectral images as in ^19^) to pinpoint areas where sampling might be most fruitful. This is likely to be of greatest importance at sites where collagen preservation is poor, and where finding a few gems amidst the dross will be most crucial. At a different scale, the speed and cost effectiveness of the technique could make it easier to address questions about inter- or intra-site variation in preservation and post-depositional processes^15,16^.

We are not suggesting that NIR spectroscopy should supplant %N or other spectroscopic techniques for addressing the question “Does this individual specimen have sufficient collagen for analysis?” However, the results presented here suggest that this tool has significant advantages over commonly employed techniques for answering the question, “Which bones at this site (or in this collection) are well preserved?” Near-infrared spectroscopy can do this form of cherry-picking quickly and inexpensively on site. This might save weeks of lab work, not to mention considerable analytical and labor costs. Most importantly, however, with such prescreening only a small number of specimens would be exposed to destructive analysis.

## Acknowledgements

This study was funded by the Max Planck Society, the Center to Advance Research and Teaching in the Social Sciences at the University of Colorado Boulder, the Office of the VCR Innovative Seed Grant Program at the University of Colorado Boulder, and the Arts and Sciences Fund for Excellence at the University of Colorado Boulder.

## Author Contributions

M.S. conceived the study. M.S., C.R., H.F., and S.T. designed the research. W.P., H.F., and S.T. provided samples for analysis. C.R. and E.S. scanned the material. M.S. and C.R. carried out the data analysis. M.S., C.R., H.F., W.P., and S.T. wrote the manuscript.

## Methods

### NIR Spectroscopy

We used 50 archaeological ground bone samples of Holocene age to create our proof of concept NIR model. All samples were scanned (50 scans per sample) while still in their glass storage vials using a fiber-optic reflectance probe attached to a LabSpec 4 NIR spectrometer with a spectral range of 350 nm to 2500 nm. Subsequent data transformations and analyses were undertaken using Unscrambler X. A Savitzky-Golay transformation (Derivative Order, 2; Polynomial Order, 3; Smoothing Points, 31) was performed to correct for additive and multiplicative effects in the spectral data^21^. Principal component analysis (PCA) was carried out to ascertain whether or not patterns relating to collagen preservation existed in the spectral dataset. After it was apparent that there were clear spectral differences relating to collagen content, the data were sorted by collagen yield and the even and odd samples were assigned to the calibration and validation datasets respectively (25 calibration, 25 validation). PLSR was then used on the calibration dataset to create a model predicting %coll^22^. The model was then applied to the validation spectra to predict %coll and assign specimens to the above or below 3% groups. After this, the same procedures were used on samples of whole bone of Holocene and Late Pleistocene age which were scanned at exposed cross-sections (Supplemental Figure 3). These scans were then coupled with scans of ground bone (48 calibration, 49 validation) in the hope of producing a PLSR model that was less sensitive to the differing geometries and particle sizes of the ground and whole bone samples^21,22^. Both models shown here use the 1695-1740 nm and 2000-2300 nm spectral ranges because previous research has shown that 1695-1740 nm, 2050 nm, 2180 nm, and 2270-2290 nm are associated with proteins including collagen^20,23^, and because a PCA loadings plot of the spectra used here confirms that these bands are associated with collagen (Figure 1c). It is worth noting, however, that it is possible to generate models with similar predictive power using data from three or fewer bands.

### Collagen Extraction

Extractions of collagen for ground bone specimens of Holocene age took place in the Archaeological Stable Isotope Lab at the University of Miami following a modified version of Longin^24^. Weighed 0.5 g aliquots of coarsely ground (0.5-1.0 mm) cortical bone were placed in 50 ml centrifuge tubes, to which 30 ml of 0.2 M HCl was added. Tubes were placed in a rotator for 24 h, at which time the degree of demineralization was assessed. Samples requiring another 24 h to demineralize had their acid refreshed at this time. After demineralization, samples were rinsed to neutral and treated with 30 ml of 0.0625 M NaOH for a period 20 h. Samples were then rinsed to neutral and gelatinized for 48 h at 90° C in 10^−3^ M HCl. The resulting gelatin was then filtered using 40 μm sterile single-use Millipore Steriflip ® vacuum filters, allowed to condense at 85° C, frozen, and then freeze-dried. Collagen yields were then determined to assess the state of sample preservation.

Extractions of collagen for whole bone specimens of Pleistocene and Holocene age took place at the Max Planck Institute for Evolutionary Anthropology in Leipzig using a modification of Method C from ^25^. About 0.5 grams of whole bone was decalcificied in 0.5 M HCl at 5°C. Acid was refreshed up to twice per week until demineralization was complete. After demineralization, samples were rinsed with ultra-pure water to neutral pH and treated with 0.1 M NaOH at room temperature for 30 minutes to remove humic acids. This was followed by a 0.5 M HCl step to remove potential contamination from modern CO2 taken up by the NaOH. Samples were rinsed to neutral pH again with ultra-pure water and gelatinized for 20 h at 75° C in 10^−3^ M HCl. The resulting gelatin was then filtered using precleaned Ezee filters (Elkay Labs UK) to remove larger particles and then ultrafiltered (precleaned Sartorius Vivapsin Turbo 15) to separate large (>30 kD) and small molecular weight fractions. The >30kD fraction was then freeze-dried for 48 hours after which collagen yields were calculated.

## Supplementary Information

**Supplementary Figure 1:**
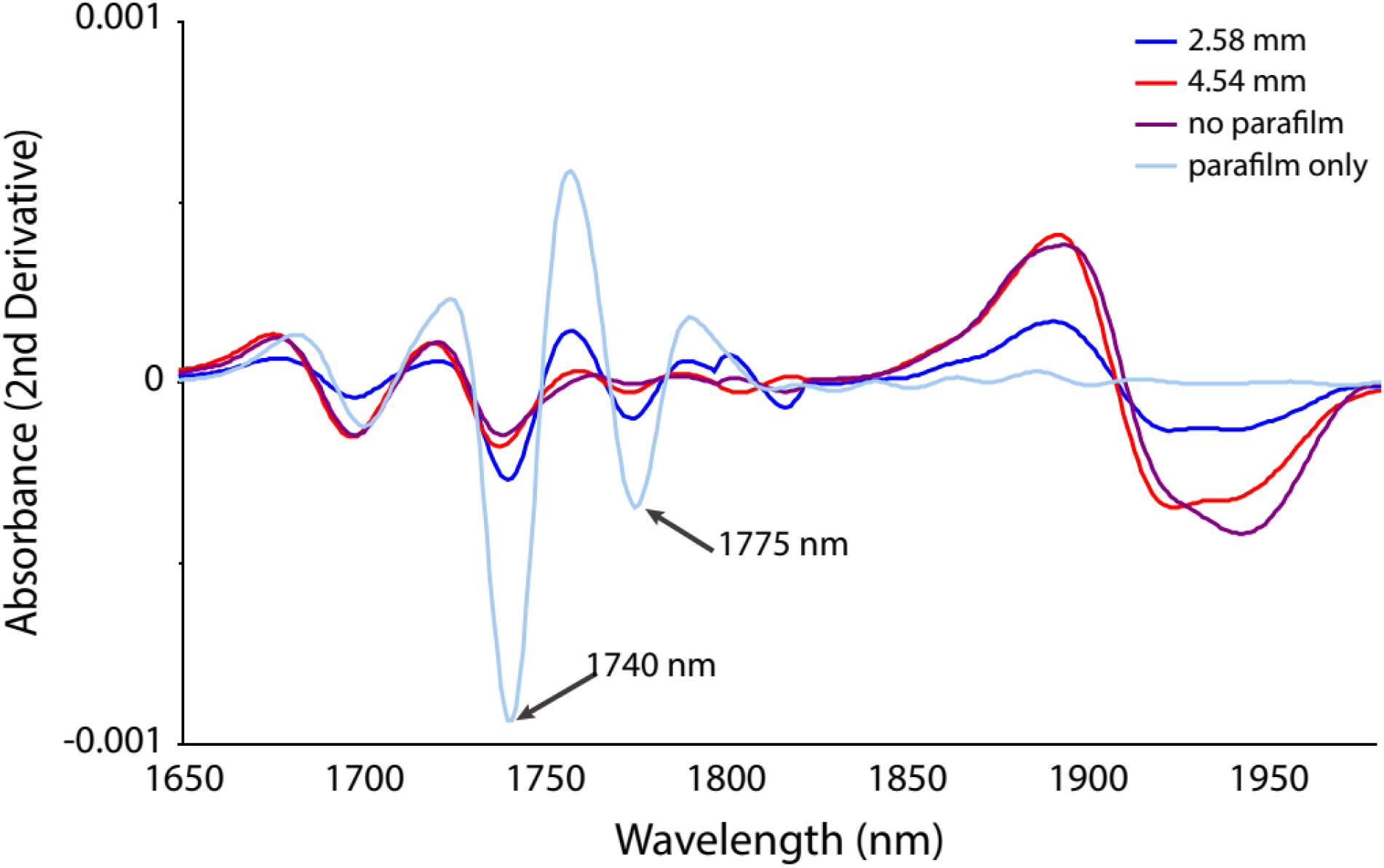
NIR penetrance in bone. Second derivative of NIR spectra of parafilm (grey), 2.58 mm of bone with parafilm beneath it (blue), 4.54 mm of bone with parafilm beneath it (red), and thick bone with no parafilm beneath it (green). Note the strongly similar spectra for the 4.58 mm bone slice and the bone not underlain by parafilm. This suggests that the NIR signal is not readily extending beyond 4.58 mm in this spectral region. In contrast, parafilm peaks at 1740 nm and 1775 nm (among others) clearly influence the spectrum of the 2.58 mm bone slice. Thus, a penetration depth of 3-4 mm for this spectral region seems reasonable and is consistent with what has been found in other mineralized tissues^1^.

**Supplementary Figure 2:**
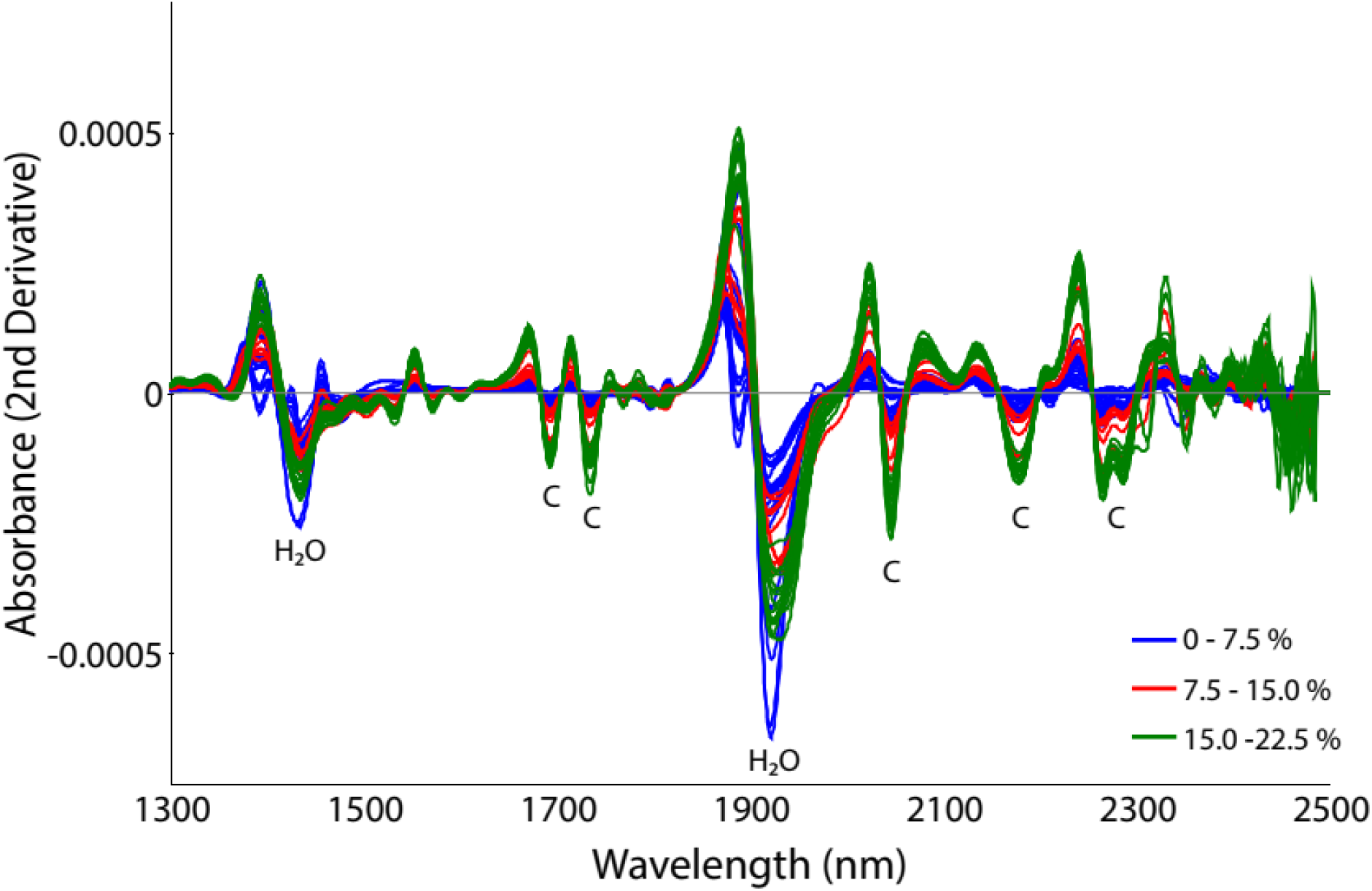
NIR spectra from ground bone samples reveal collagen content. A) Second derivative of NIR spectra from all 50 ground bone specimens in this study. Green spectra are high %coll specimens (15.0-22.5%), red spectra have less collagen (7.5-15%), while blue spectra have the lowest %coll (0.0-7.5%). Bands/regions that show strong patterning with %coll are labeled C. Water bands (labeled H_2_0) show less consistent patterning with collagen and were not used for PLSR.

**Supplementary Figure 3:**
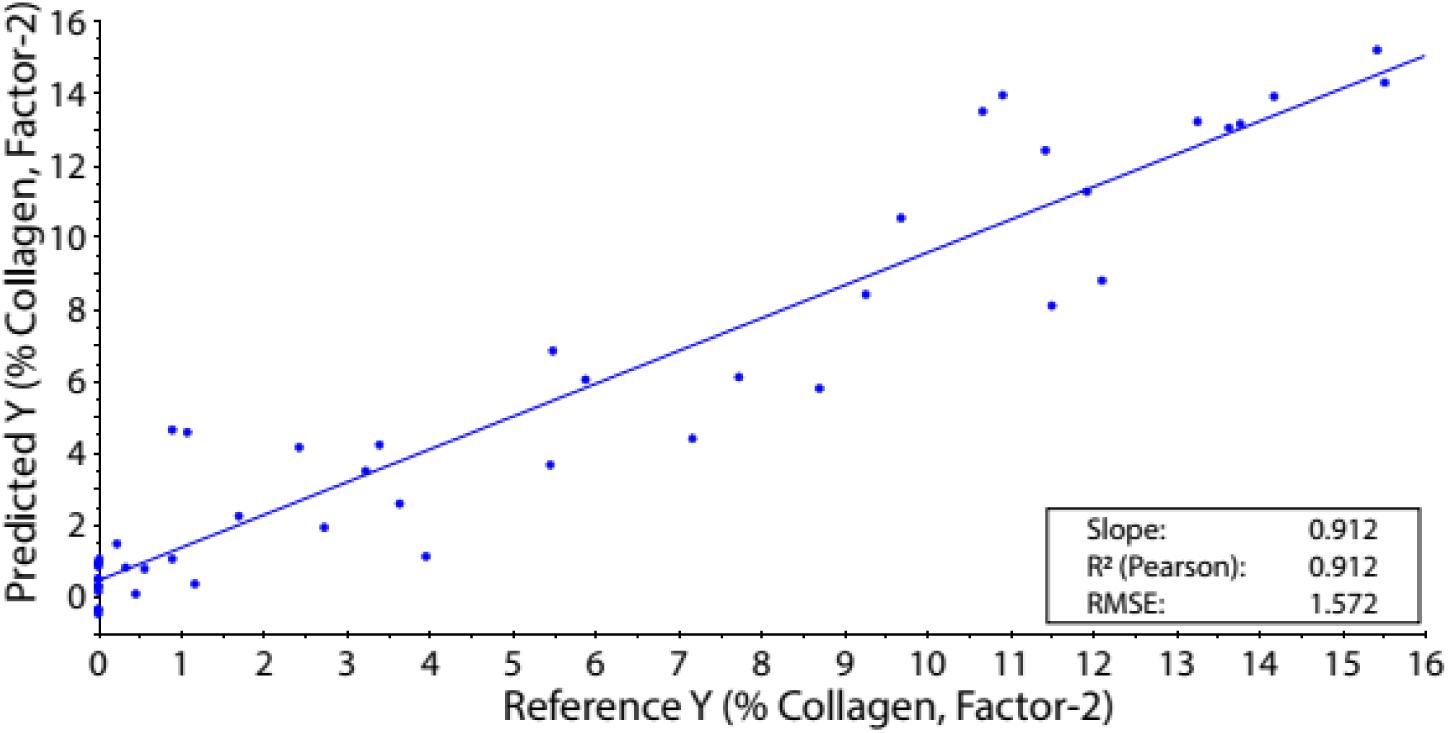
Predicting collagen preservation from NIR spectra of bone cross sections. A) Results of PLSR showing predicted versus actual %coll values for 47 whole bone samples ranging in age from 2,000 to about 45,000 years old. Bones were scanned along previously and naturally exposed cross sections. The model’s performance is good (R2 = 0.91; RMSEC = 1.6; leave-one-out cross validation R2 = 0.90; Root Mean Square Error of Cross Validation = 1.7), although not as strong as the ground bone model (Figure 1a). This was expected. Collagen yield had been determined for both the ground and whole bone specimens (for radiocarbon and isotopic paleodietary studies) before spectra were acquired. Given the relative homogeneity of ground bone, the scans taken were almost certainly representative of the subsamples on which collagen determinations were made. Whole bone could not be scanned at the locations where %coll was determined as the sample is transformed during that process. Given that collagen preservation is heterogeneous within whole bone^2^, it is likely that, at least in some cases, areas were scanned that had different %coll than the locations where collagen determinations were made. Thus, the ground bone and ground/whole bone models were thought to be better proofs of concept.

**Supplementary Table 1.** Specimen IDs, areas of origin, approximate ages, and collagen yields (%wt) for specimens included in this study. Italicized specimens were analyzed as ground bone.

| **Specimen** | **Country/Territory** | **Thousands of Years**<br>**BP (approx)** | **Collagen**<br>**Yield (wt%)** |
| --- | --- | --- | --- |
| R1(1) | Italy | 45 | 0 |
| R2(1) | Italy | 45 | 0.2 |
| R3(1) | Italy | 45 | 0.5 |
| R4 | Italy | 45 | 0 |
| R5 | Italy | 45 | 0.3 |
| R6 | Italy | 45 | 1.1 |
| R7 | Italy | 45 | 0 |
| R8 | Italy | 45 | 0.6 |
| R9 | Mongolia | 45 | 11.9 |
| R10 | Serbia | 25-45 | 7.2 |
| R11 | Serbia | 25-45 | 13.8 |
| R12 | Serbia | 25-45 | 4 |
| R13 | Serbia | 25-45 | 9.3 |
| R14 | Serbia | 25-45 | 9.7 |
| R28 | Gibraltar | 45 | 0.9 |
| R29 | Italy | 25 | 0.9 |
| R30 | Spain | 50-30 | 5.5 |
| R31 | Spain | 50-30 | 5.9 |
| R32 | Spain | 50-30 | 2.7 |
| R33 | Spain | 50-30 | 1.7 |
| R34 | Spain | 50-30 | 3.2 |
| R35 | Spain | 50-30 | 1.2 |
| R36 | Spain | 50-30 | 0 |
| R37 | Italy | 50-40 | 0 |
| R38 | Italy | 50-40 | 0 |
| R39 | Italy | 50-40 | 0 |
| R40 | Italy | 50-40 | 0 |
| R41 | Italy | 50-40 | 0 |
| R42 | Hungary | 14-13 | 7.7 |
| R43 | Hungary | 14-13 | 3.6 |
| R44 | Hungary | 14-13 | 5.5 |
| R45 | Hungary | 14-13 | 2.4 |
| R46 | Gibraltar | 45 | 0 |
| R47 | Gibraltar | 45 | 0 |
| R48 | Gibraltar | 45 | 0 |
| R49 | Canada | 2 | 13.3 |
| R50 | Canada | 2 | 10.9 |
| R51 | Canada | 2 | 15.4 |
| R52 | Canada | 2 | 12.1 |
| R53 | Canada | 2 | 15.5 |
| R54 | Canada | 2 | 14.2 |
| R55 | Canada | 2 | 13.6 |
| R56 | Mongolia | 33 | 11.4 |
| R57 | Mongolia | 33 | 10.7 |
| R1 | Austria | >50 | 11.5 |
| R2 | Czech Republic | 26 | 8.7 |
| R3 | Czech Republic | 26 | 3.4 |
| *A-110* | *Puerto Rico* | *1* | *9.4* |
| *A-13* | *Puerto Rico* | *1* | *4.2* |
| *A-18* | *Puerto Rico* | *1* | *4.4* |
| *A-31* | *Chile* | *2* | *0.9* |
| *A-45* | *Puerto Rico* | *1* | *6.1* |
| *A-47* | *Puerto Rico* | *1* | *5.9* |
| *A-52* | *Puerto Rico* | *1* | *6.7* |
| *A-53* | *Puerto Rico* | *1.5* | *6.4* |
| *A-56* | *Puerto Rico* | *1.5* | *6* |
| *A-58* | *Chile* | *1.5* | *1.3* |
| *A-60* | *Puerto Rico* | *1.5* | *2.4* |
| *A-63* | *Puerto Rico* | *1* | *9.7* |
| *A-83* | *Puerto Rico* | *1* | *7.9* |
| *C-18* | *Puerto Rico* | *1.5* | *9.9* |
| *F-18* | *Puerto Rico* | *1* | *1* |
| *F-22* | *Puerto Rico* | *1* | *5.1* |
| *F-28* | *Puerto Rico* | *1* | *5.9* |
| *F-3* | *Puerto Rico* | *1* | *4.2* |
| *F-32* | *Puerto Rico* | *1* | *2.2* |
| *F-35* | *Puerto Rico* | *1* | *7.1* |
| *F-42* | *Puerto Rico* | *1* | *6.9* |
| *F-5* | *Puerto Rico* | *1* | *4.2* |
| *F-55* | *Puerto Rico* | *1* | *5.8* |
| *F-63* | *Iraq* | *4* | *1.1* |
| *F-71* | *Chile* | *2.5* | *17.8* |
| *F-76* | *Chile* | *2* | *21.7* |
| *F-92* | *Chile* | *1* | *11.8* |
| *G-20* | *Chile* | *1* | *12* |
| *G-21* | *Chile* | *1* | *16.1* |
| *G-43* | *Chile* | *1* | *0.9* |
| *G-46* | *Chile* | *1* | *19.2* |
| *G-59* | *Kenya* | *0.5* | *3.5* |
| *G-8* | *Chile* | *1* | *19.6* |
| *H-101* | *Chile* | *1* | *20.4* |
| *H-104* | *Chile* | *1* | *22.1* |
| *H-105* | *Chile* | *1* | *4.8* |
| *H-112* | *Chile* | *1* | *1.5* |
| *H-13* | *Chile* | *1* | *18.4* |
| *H-46* | *Chile* | *2* | *10.7* |
| *H-78* | *Chile* | *1* | *15.6* |
| *H-80* | *Chile* | *1* | *8.9* |
| *H-9* | *Chile* | *1* | *21.4* |
| *H-90* | *Chile* | *0.5* | *21.5* |
| *H-95* | *Chile* | *0.5* | *19.2* |
| *I-110* | *Chile* | *2.5* | *21.1* |
| *I-2* | *Chile* | *1.5* | *2.9* |
| *I-25* | *Chile* | *1.5* | *15.9* |
| *I-48* | *Puerto Rico* | *1.5* | *1.7* |
| *J-2* | *Chile* | *2.5* | *14.1* |
| *J-3* | *Chile* | *2.5* | *17.8* |

